# Modeling the Synergetic Dynamics of B cells and TFH cells in Germinal Center Reactions

**DOI:** 10.1101/2025.11.27.691027

**Authors:** Andrew G. T. Pyo, Julia Merkenschlager, Quan Pham, Benjamin H. Good, Michel C. Nussenzweig, Ned S. Wingreen

**Affiliations:** Department of Applied Physics, Stanford University, Stanford, CA; Department of Physics, Princeton University, Princeton, NJ; Department of Immunology, Harvard Medical School, Boston, MA; Laboratory of Molecular Immunology, The Rockefeller University, New York, NY; Department of Biology, Stanford University, Stanford, CA; Biohub, San Francisco, CA; Howard Hughes Medical Institute, The Rockefeller University, New York, NY; Lewis-Sigler Institute for Integrative Genomics, Princeton University, Princeton, NJ; Department of Molecular Biology, Princeton University, Princeton, NJ

## Abstract

B cells producing high-affinity antibodies arise through affinity maturation within germinal centers (GCs), where selection is driven by T follicular helper (T_FH_) cells. Recent studies have shown that, like GC B cells, T_FH_ cells also undergo antigen-dependent selection, with competition among T_FH_ clones dictated by their ability to recognize and stimulate B cells. This sensitivity-dependent selection process leads to dynamic remodeling of the T_FH_ repertoire over time. Despite the essential role of T_FH_ cells in B cell selection, the functional consequences of the time evolution of the T_FH_ cell population remains poorly understood. To address this gap, we developed a population dynamics model that explicitly incorporates key T_FH_ cell properties and dynamics. Our analysis predicts that dynamic feedback between B and T_FH_ cell populations provides robust homeostatic regulation of their numbers in the GC, yielding a stable lymphocyte ratio that we verify experimentally. Moreover, our model predicts that T_FH_ clone sensitivity dictates distinct evolutionary strategies during affinity maturation, with low-sensitivity T_FH_ cells accelerating affinity gain at the expense of B cell diversity, while high-sensitivity T_FH_ cells slow affinity maturation but preserve a broader B cell repertoire. These findings highlight the importance of co-regulation between T_FH_ and B cells and suggest that reciprocal stimulation allows the immune system to tune the tradeoff between the speed of affinity gain and the breadth of B cell diversity—a principle that may extend to other adaptive systems.

**Significance Statement:** Effector B cells that secrete high-affinity antibodies and form immunological memory are essential for humoral immunity and arise from germinal center (GC) reactions. Within GCs, B cells undergo an accelerated version of Darwinian evolution to enhance antibody affinity. This process is orchestrated by T follicular helper (T_FH_) cells which provide stimulatory signals to selected B cells and undergo their own antigen-driven selection. To investigate this co-evolutionary process, we developed a tractable population-level model of the GC reaction. Our analysis reveals that the reciprocal stimulation of B and T_FH_ cells provides a robust mechanism for regulating the B:T_FH_ ratio and tuning the tradeoff between the speed of affinity maturation and the diversity of the antibody response.

## Introduction

Humoral immunity including an effective response to vaccinations relies on the timely production of high-affinity, antibody-producing B cells from germinal centers (GCs) (1). GCs are specialized microenvironments where B cells expressing high-affinity antibodies are evolved from low-affinity precursors (2, 3). This affinity maturation process involves iterative cycles of divisions accompanied by somatic hypermutations (SHM) (4). However, because SHM is largely random, the great majority of mutations are deleterious (i.e., affinity-decreasing), rather than beneficial (i.e., affinity-enhancing), necessitating a stringent selection process to achieve affinity maturation (5). This selection is mediated by T follicular helper (T_FH_) cells. B cells compete to capture antigen from follicular dendritic cells (FDCs) (6), and during stochastic encounters with T_FH_ cells, the B cells’ capacity to bind, internalize, and present antigen dictates the level of T_FH_-derived help they receive, which determines their proliferative capacity (7, 8). B cells that fail to engage with cognate T_FH_ cells undergo negative selection by apoptosis and are removed from the GC reaction. In contrast, positively selected B cells can either leave the GC by differentiating into plasma or memory cells, or continue iterative cycling in the GC (9, 10).

The degree of T_FH_ cell help is therefore a key determinant of GC dynamics. It is well established that T_FH_ cell numbers must be limiting in order to maintain appropriate selection pressure on B cells. If T_FH_ cells are too abundant, selection becomes permissive, whereas too few renders it overly restrictive, in both cases leading to abnormal GC outcome (1, 11, 12). Consistent with this notion, the number of B cells strongly correlate to the number of T_FH_ cells within GCs, and remains remarkably consistent across diverse antigens and individuals (13–17) even under varying levels of available antigen and co-stimulatory ligands (18, 19). However, the mechanisms that maintain the proportion of T_FH_ cells in the GC, and how this balance influences B cell affinity maturation remain unclear.

Regulation of the number of GC lymphocytes may in part arise from the complex clonal and functional dynamics that T_FH_ cells undergo throughout a primary GC reaction (19–21). In particular, preferential expansion of T_FH_ cell clones bearing T cell receptors (TCRs) with higher sensitivity to peptide–major histocompatibility complexes (pMHCs) has been observed (20). Within the GC, T_FH_ cells interact with antigen presented both by B cells and by FDCs (1, 11). Thus, variation in TCR fitness, together with interactions mediated by co-stimulatory receptors such as ICOSL and CD40L, drive competition among T_FH_ clones through differential proliferation and survival. This competition leads to dynamic changes in both T_FH_ cell abundance and TCR clonotype composition, which shape ongoing GC B cell selection. In late stage GCs, the T cell compartment undergoes further remodeling, with regulatory T follicular cells dominating over conventional T_FH_ as the GC collapses (21–23). These observations underscore the intricate interplay between lymphocytes in the GC, raising the question of how the dynamic behavior and quality of T_FH_ cells influence the efficiency and outcome of B cell affinity maturation. Previous theoretical approaches, from probabilistic models to detailed agent-based simulations, have provided insight into various aspects of the GC reaction (24–30). However, despite the pivotal role of T_FH_ cells in B cell affinity maturation, models explicitly examining how T_FH_ cell properties and dynamics influence the GC reaction are lacking.

To address this question, we developed a population-level model of the GC reaction that explicitly incorporates the coupling of T_FH_ cell dynamics to B cell affinity maturation. Using our framework, we find that reciprocal interactions between B cells and T_FH_ cells give rise to a “homeostatic” mechanism that maintains a remarkably consistent B:T_FH_ ratio, regardless of variations in antigen availability, T_FH_ cell sensitivity (i.e. propensity to provide help), or the affinity of B cell receptors. Consistent with this prediction, *in vivo* measurements confirm that the B:T_FH_ ratio is maintained across time and different antigen availabilities. Moreover, we found that T_FH_ cell sensitivity strongly modulates both the breadth and depth of the GC B cell repertoire: high-sensitivity T_FH_ cells promote the diversification of the GC B cell repertoire, but at the cost of slower affinity maturation and specificity toward dominant epitopes. Together, these results reveal a key regulatory mechanism, enabled by dynamic T_FH_–B cell interactions, that underpins effective GC function and affinity maturation.

## Results

### Population dynamics model of the GC reaction

To investigate the influence of T_FH_ cells on B cell affinity maturation, we developed a population dynamics model that explicitly incorporates feedback between lymphocytes. The model describes B cell affinity maturation as an evolutionary hill-climbing process, wherein mutation and selection cause GC B cells to progressively improve their affinity. These GC B cells are categorized into discrete affinity classes, labeled by *n*, where each step *n* → *n+*1 corresponds to an increase in the binding energy between the antigen and the B cell receptor (BCR) by an incremental amount *ɛ* (31). Upon division, each daughter cell acquires mutations with probability *p_n_* (27), where we account for neutral, beneficial, and deleterious mutations. A beneficial mutation increases the B cell’s affinity class (*n* → *n+*1), while a deleterious mutation decreases it (*n* → *n*−1).

B cell selection is determined by the effective B cell division rate, which depends on the ability of the B cell to capture antigen (Figure 1, i), and subsequent encounter and help from T_FH_ cells (Figure 1, ii). When antigen availability is the limiting resource, the selection strength for a B cell in affinity class *n* (denoted *B_n_*) interacting with T_FH_ cells of clone *σ* (denoted *T_ᓂ_*) is given by

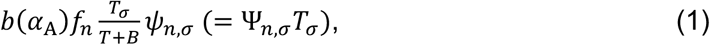

**Figure 1.**
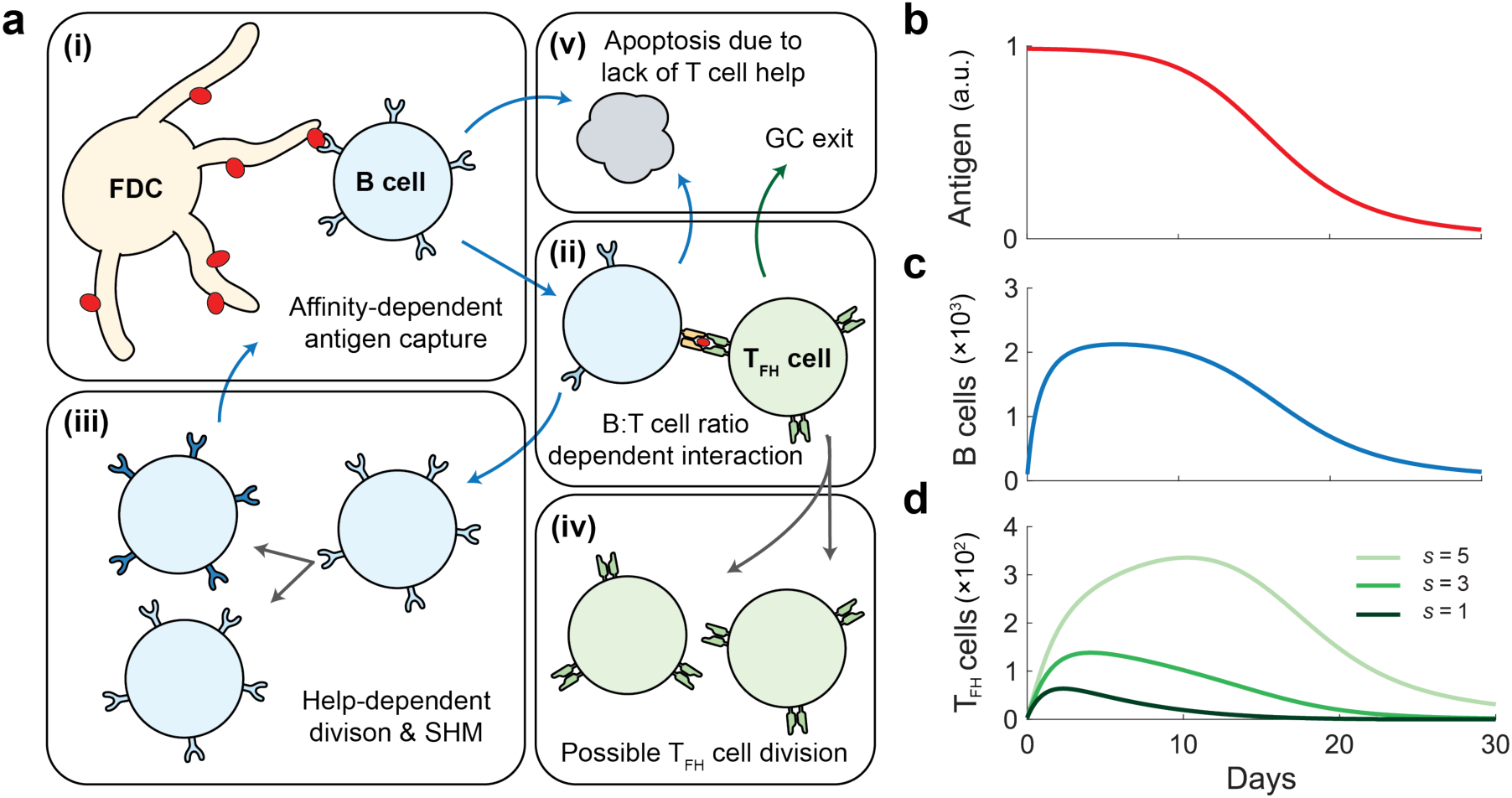
B cell affinity maturation model. **a)** Schematic of the model framework. B cells capture antigen from FDCs in an affinity-dependent manner (i). T_FH_ cells interact with B cells based on their relative proportions in the GC and provide help signals proportional to the amount of presented antigen (ii). B cells undergo rounds of division accompanied by SHM in proportion to the help signal (iii), while T_FH_ cells may divide following productive interactions with B cells (iv). B cells that fail to receive sufficient help undergo apoptosis, and T_FH_ cells exit the GC at a fixed rate (v). **b–d)** Example of model dynamics for antigen availability decaying over time **(b)**. B cell population initially expands and subsequently contracts as antigen becomes limiting **(c)**. Clonal dynamics of T_FH_ cells show that high-sensitivity clones (*s* = 5, light green) outcompete lower-sensitivity clones (*s* = 1, 3, darker greens) **(d)**.

where *b*(*⍺*_A_) is the overall fraction of B cells that have captured antigen which depends on the rate of antigen influx *⍺*_A_, *f_n_* quantifies the relative antigen-capture efficiency of affinity class *n*, *B* = ∑*_n_ B_n_* is the total number of B cells, *T* = ∑*_ᓂ_ T_ᓂ_* is the total number of T_FH_ cells, and *ψ_n_*,*_ᓂ_* represents the sensitivity-dependent propensity of T_FH_ clone *σ* to provide help to B cells of affinity class *n*. Here, T_FH_ cell sensitivity reflects underlying factors such as TCR affinity and the strength of co-stimulatory interactions. In Equation 1, the first factor (*bf_n_*) describes antigen presentation across B cell affinity classes, the second factor 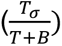 represents the likelihood of B cells in affinity class *n* encountering a particular T_FH_ cell clone, and the last factor (*ψ_n_*,*_ᓂ_*) quantifies the amount of help signal received.

B cell proliferation depends on this T cell help (Figure 1, iii), while cells that fail to receive help undergo apoptosis at a fixed rate (Figure 1, v). These processes are described by

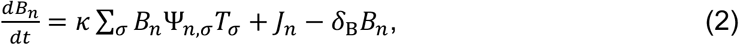

where ***κ*** is the maximum effective rate of B cell division, *J_n_* represents the flux due to mutations (see SI), and *δ*_*_ is the B cell death rate. Similarly, T_FH_ cells may proliferate following productive interactions with B cells (Figure 1, iv), quantified by the relative probability factor *γ*, and exit the GC at a fixed rate *δ*_+_ (Figure 1, v). Their dynamics are given by

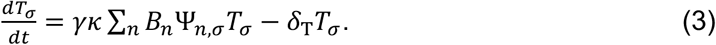

Together, the ordinary differential equations (ODEs) shown in Equations 2 & 3 describe the co-evolution of B cell and T_FH_ cell populations throughout the course of a GC reaction. For deterministically decaying of antigen availability (Figure 1b), our model recapitulates the corresponding rise and fall of the B cell population (Figure 1c), and the sensitivity-dependent competition among T_FH_ cell clones (Figure 1d). Detailed derivations and parameter values can be found in the SI.

### Stable B:T_FH_ cell ratio emerges from GC dynamics

The reciprocal interaction between B and T_FH_ cells provides a potential mechanism by which lymphocyte populations can be regulated in GC reaction. To characterize the resulting population-level behavior of lymphocytes in our model, we analyzed the system’s steady-state dynamics by fixing the rate of antigen influx (*⍺*_A_) to a constant value, and examined how the resulting equilibrium populations depended on model parameters. Under these conditions, the lymphocyte dynamics (Equations 2 & 3) admits a single, linearly-stable fixed-point (see SI). At steady state, the total number of lymphocytes is given by

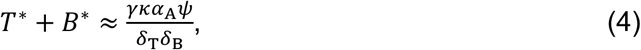

which depends on T_FH_ cells sensitivity through *ψ*, and on antigen availability through *⍺*_A_, the rate of antigen influx. The approximation in Equation 4 was obtained assuming a simplified case with identical B cells (i.e., no affinity classes or mutation flux) interacting with a monoclonal population of T_FH_ cells. The steady-state ratio of B to T_FH_ cells is

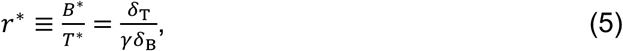

Notably, the ratio *r*^∗^ is independent of antigen availability or T_FH_ cell sensitivity.

As shown in Figure 2a, simulations of B cell affinity maturation (Equations 2 & 3) with monoclonal T_FH_ cells under conditions of higher antigen availability (larger circles) and increased T_FH_ cell sensitivity (lighter green) led to larger populations of B cells and T_FH_ cells within the GC, as expected from Equation 4. Despite these substantial changes in cell numbers, the B:T_FH_ ratio remained constant, consistent with Equation 5 (Figure 2a, dotted line). These results indicate that reciprocal coupling of B cell and T_FH_ cell proliferation could provide a robust mechanism to maintain a stable B:T_FH_ cell ratio in the GC.

**Figure 2.**
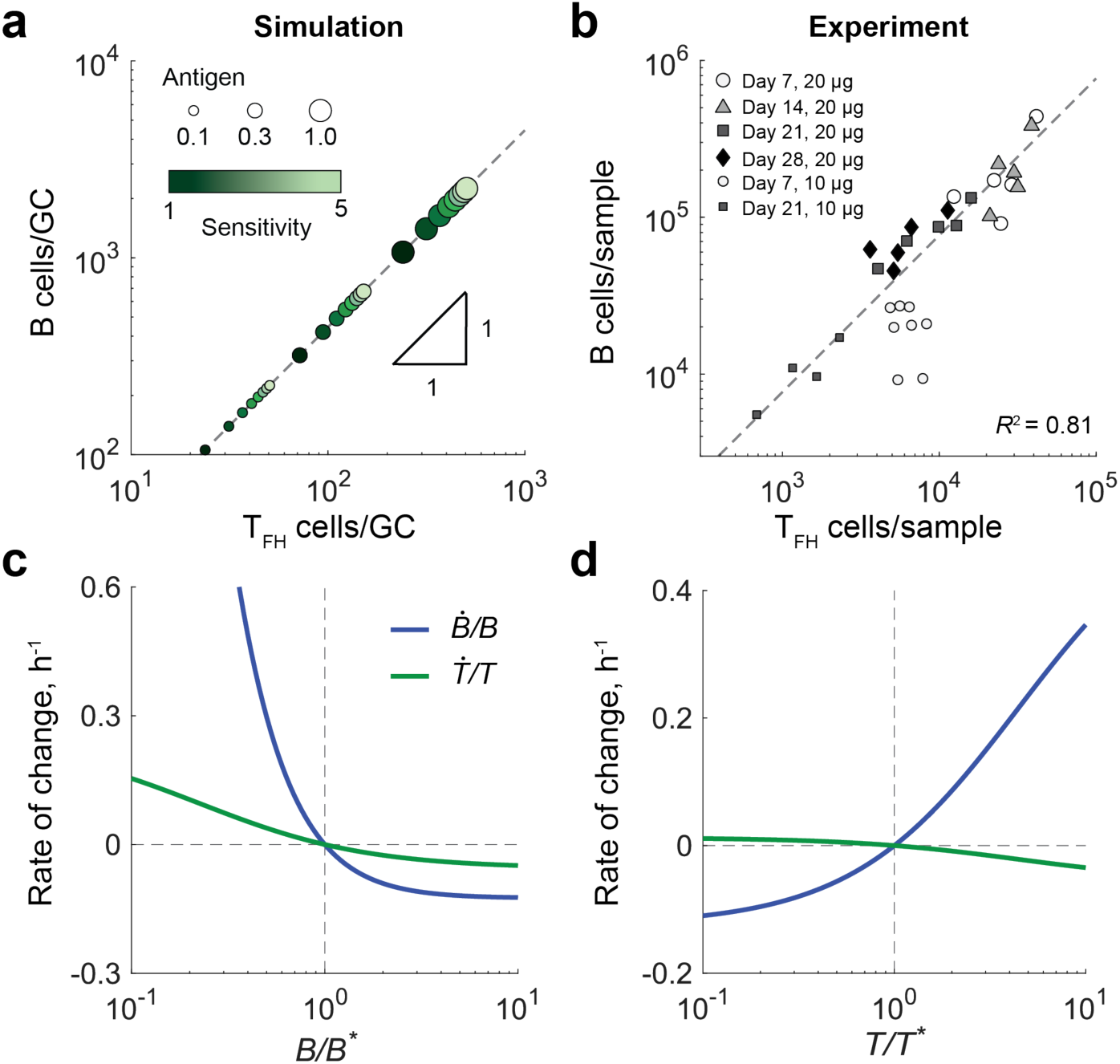
Homeostatic regulation of the B:T_FH_ ratio in the GC. **a)** Influence of antigen availability and T_FH_ cell sensitivity on GC cell numbers. Steady-state numbers of B cells and T_FH_ cells are shown across conditions of high antigen availability (largest circles) to low antigen availability (smallest circles). Antigen levels are reported in dimensionless units, normalized by *⍺*_A_/(1000*δ*_*_). T_FH_ cell sensitivity is indicated by shading, ranging from low sensitivity (s = 1, dark green) to high sensitivity (s = 5, light green). Constant ratio line is shown as a gray dashed line. **b)** Experimental measurement of GC B and T_FH_ cell numbers. Data re shown for mice receiving a 20µg bolus (large symbols) or a 10µg bolus (small symbols). A constant B:T_FH_ ratio fit yielded *r* = 7.7 ± 0.9 (gray dashed line). **c, d)** Per capita growth rates of B cells (blue) and T_FH_ cells (green) following perturbations in the B cell population **(c)** or the T_FH_ cell population **(d)**. Note that the rate of change in the B:T_FH_ ratio is given by 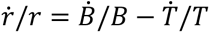.

### Experimental validation of a stable GC B:T_FH_ ratio

Our model predicts that the reciprocal interaction between B and T_FH_ cells leads to a stable B:T_FH_ ratio even when total cell number varies due to antigen availability or T_FH_ cell sensitivity. To test this prediction experimentally, we immunized wild-type (wt) mice with a bolus of 10μg or 20μg of 4-hydroxy-3-nitrophenylacetyl-ovalbumin (NP-OVA) and then quantified the relative abundance of GC B and T_FH_ cell populations throughout the course of GC reactions.

As shown in Figure 2b, GC B cell numbers peak around day 7, after which both GC B and T_FH_ cell populations progressively decline. Notably, larger antigen boluses (Figure 2b, large symbols) induce GC reactions that sustain lymphocyte populations over an order of magnitude higher than those observed with smaller boluses (Figure 2b, small symbols). Despite the substantial changes in the absolute numbers of GC B and T_FH_ cells over time and bolus size, the B:T_FH_ cell ratio remained remarkably stable over time (Figure 2b, gray dotted line), with an average ratio of *r* = 7.7 ± 0.9.

### B:T_FH_ ratio in the GC is homeostatically maintained

After observing that the B:T_FH_ ratio is sustained overtime and demonstrating that reciprocal interactions can account for this behavior, we next asked how this ratio is robustly maintained. To address this, we used the reduced version of the model in which identical B cells (neglecting affinity classes) are selected by a monoclonal T_FH_ population. At steady state, we examined the changes in the net rates of B cell and T_FH_ cell proliferation when either population was perturbed from its steady-state number.

When the B cell population is perturbed, its net growth counter acts the change: B cell numbers increase if they fall below the equilibrium value and decrease if they exceed it (Figure 2c). This occurs because, according to Equation 1, a smaller proportion of B cells means each B cell has a higher probability of encountering a T_FH_ cell, while a larger B cell proportion reduces this probability. For the same reason, perturbations in the T_FH_ cell population also lead to compensatory changes in B cells: if the T_FH_ cell population falls below its steady-state level, B cell numbers decrease, whereas if T_FH_ cells exceed their equilibrium, B cell numbers increase (Figure 2d). In both cases, changes in the T_FH_ cell population occur more slowly, reflecting its inherently slower dynamics (Figure 2c & d, green curves). Importantly, the rate of change *ṙ*/*r* of the B:T_FH_ ratio is proportional to the difference in per capita growth rates of B and T_FH_ cells (i.e. *ṙ*/*r* = *Ḃ*/*B* − *Ṫ*/*T*). Thus, in both cases, the B:T_FH_ ratio is homeostatically regulated by the coupled dynamics of B cells and T_FH_ cells, resulting in the maintenance of a constant B:T_FH_ ratio.

### High-sensitivity T_FH_ cells slow down affinity maturation

The fact that high T_FH_ cell sensitivity does not affect the overall B:T_FH_ cell ratio in the GC raises the question of whether T_FH_ cells influence B cell affinity maturation. Specifically, we examined how T_FH_ cell sensitivity impacts the rate of B cell affinity increases, used here as a proxy for the efficiency of affinity maturation. To address this, we simulated the dynamics described by Equations 2 & 3, initiating the GC reaction with 100 low-affinity B cells and six T_FH_ cells of identical sensitivity. After allowing the system to reach steady-state population sizes, we calculated the rate of increase in the mean B cell binding affinity.

We found that increasing T_FH_ cell sensitivity led to a slower rate of affinity maturation (Figure 3a). This occurred despite high-sensitivity T_FH_ cells supporting a larger overall B cell population (Figure 2a), which would typically accelerate the speed of adaptation in evolving populations (32, 33). The slowdown arose because high-sensitivity T_FH_ cells reduced the effective selection pressure on B cells: B cells expressing high-affinity antibodies experienced a smaller fitness advantage than when interacting with low-sensitivity T_FH_ cells (Figure 3b). Mechanistically, high-sensitivity T_FH_ cells, while more likely to provide help to individual B cells (Figure 3c, green line), sustain a larger B cell population, increasing competition for limited antigen (Equation 1, first factor). Resulting antigen scarcity limits the extent to which B cells can benefit from the affinity of the antibodies they express. Even B cells expressing high-affinity BCRs capture only slightly more antigen than those with lower-affinity BCRs, narrowing differences between clones and slowing the rate of affinity maturation.

**Figure 3.**
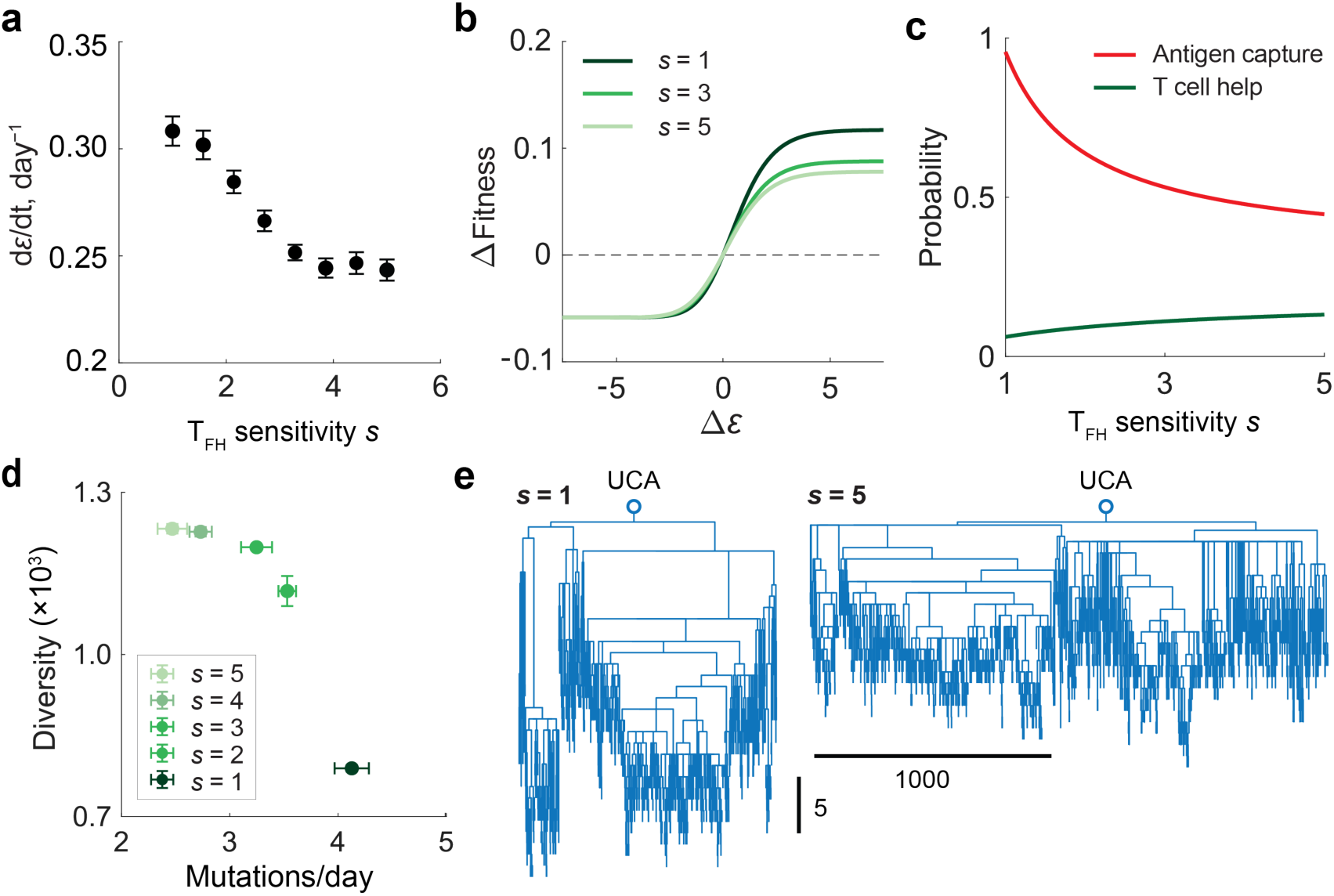
T_FH_ cell sensitivity tunes the affinity-diversity tradeoff of the GC reaction. **a)** Rate of affinity increase, quantified by the change in average B cell binding energy *ɛ* over time. Fitness difference relative to the mean B cell fitness across affinity classes (ΔFitness = *Ψ*(*ɛ*)*/κ*) for T_FH_ cells of varying sensitivities; lighter green indicates higher T_FH_ sensitivity. **b)** Average probability that a B cell captures antigen (red) and probability of obtaining T_FH_ help (green) for different T_FH_ sensitivities. **d)** Diversity of GC B cells, measured as the inverse Simpson index, and the average number of mutations acquired per day for varying T_FH_ sensitivities. Error bars indicate the standard error across five simulations. **e)** Phylogenetic trees at day 5 for B cells originating from a unmutated common ancestor (UCA) under selection from low-sensitivity (*s* = 1, left) or high-sensitivity (s = 5, right) T_FH_ cells. Tree depth indicates the number of accumulated mutations, while the width reflects the number of surviving lineages. The phylogenetic trees were generated based on the pairwise mutational distance matric using the *seqneighjoin* function implemented in MATLAB R2025a. Panels (a–c) were obtained from ODE simulations, and (d, e) from agent-based simulations. A steady antigen supply rate (*⍺*_A_ = 125 h^−1^) was used for all simulations.

### T_FH_ cell sensitivity tunes the diversity of GC B cells

Having found that high-sensitivity T_FH_ cells impose weaker selective pressure on B cells, we next asked whether this reduced selection influences the diversity of GC B cells. As the ODE model (Equations 2 & 3) only tracks the corresponding antibody affinity and not the lineage of B cells, we developed a hybrid model that combines the ODE-based population dynamics with agent-based tracking of stochastic B cell mutation histories. This framework allowed us to reconstruct the full lineage relationships among B cells (see SI).

Using this hybrid model, we again considered the scenario with steady antigen supply and monoclonal T_FH_ cell population. Precursor B cells were set to be identical, so that all diversifications arose from mutations during GC selection. From the simulations, we quantified GC B cell diversity using the inverse Simpson index and calculated the average number of mutations accumulated by surviving B cells. As shown in Figure 3d, GCs supported by high-sensitivity T_FH_ cells exhibited slower accumulation of mutations, but greater overall diversity compared to those supported by low-sensitivity T_FH_ cells. Consistently, phylogenetic trees of surviving B cells at day 5 revealed that low-sensitivity T_FH_ cells produced deeper but narrower trees, whereas high-sensitivity T_FH_ cells generated shallower trees with broader branches (Figure 3e). Together, these results indicate that T_FH_ cell sensitivity modulates the balance between selection strength and clonal diversity: high-sensitivity T_FH_ cells promote broader but less affinity-focused B cell repertoires.

### Role of B cell survival in the GC reaction

So far, we have examined how T_FH_ cell sensitivity, defined by TCR affinity for cognate peptide and the propensity to provide help, affects B cell affinity maturation. In addition to delivering help through cognate interactions, T_FH_ cells also promote B cell survival by secreting cytokines such as IL-4 and IL-21 (22, 34–37). To assess the impact on affinity maturation of this more indiscriminate support, we examined how changes in the B cell apoptosis rate influence GC dynamics. Because it remains unclear whether high-sensitivity T_FH_ cells also produce more cytokines, we held T_FH_ cell sensitivity fixed and varied the B cell lifetime in the GC as a proxy for differential cytokine support.

Under steady antigen supply (fixed *⍺*_A_) with mutations and a monoclonal T_FH_ cell population, increasing B cell lifetime led to a pronounced expansion of the steady-state B cell population, while the T_FH_ cell population remained largely unchanged (Figure 4a). This resulted in an elevated B:T_FH_ ratio (Figure 4a, inset). To determine how such shifts in population composition influence affinity maturation, we next examined the evolution of B cell affinity and diversity. Extending B cell lifetime slowed the rate of affinity maturation (Figure 4b, black) while increasing B cell diversity (Figure 4b, blue), mirroring the speed–diversity trade-off observed previously when T_FH_ cell sensitivity was varied. These findings suggest that cytokine-mediated modulation of B cell survival can shape GC outcomes by tuning the balance between the speed and breadth of the antibody response.

**Figure 4.**
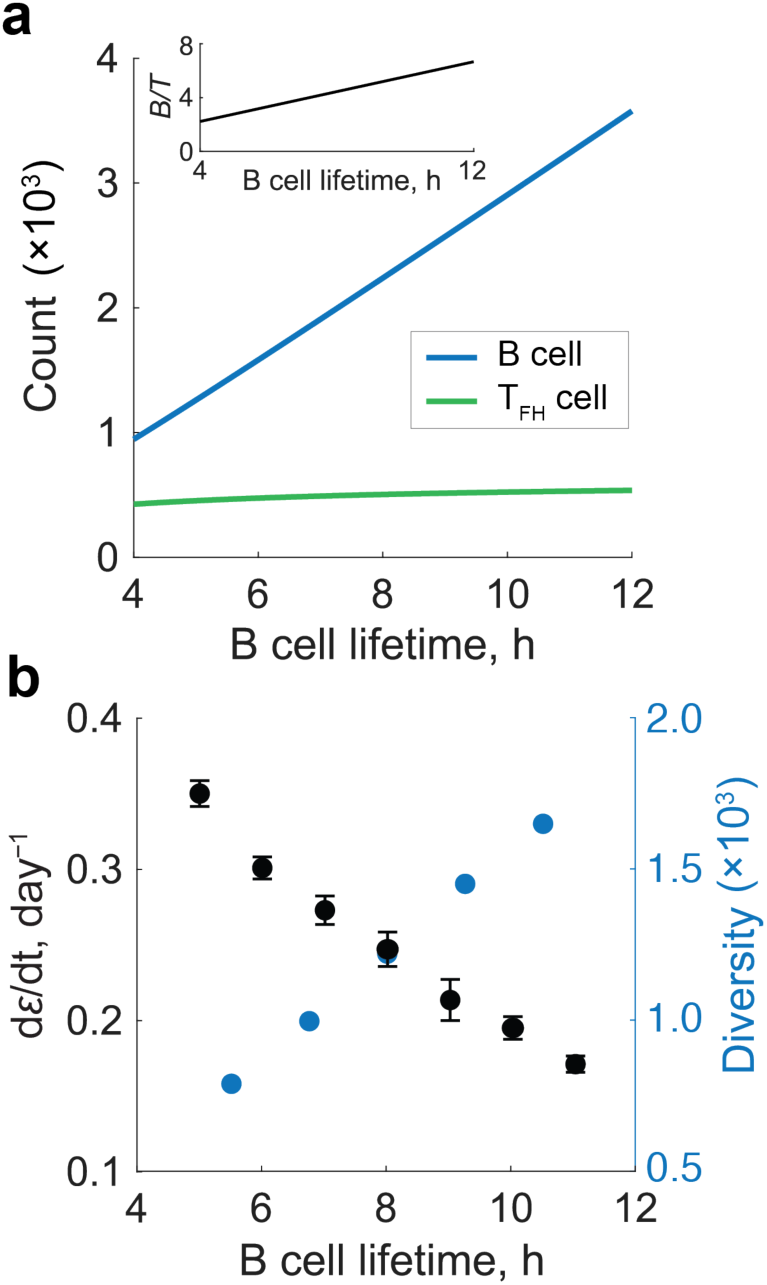
Influence of B cell lifetime on the GC reaction. **a)** Steady-state numbers of B cells (blue) and T_FH_ cells (green) for varying B cell lifetimes. The corresponding B:T_FH_ ratio is shown in the inset. **b)** Rate of affinity increase (black) and steady-state B cell diversity (blue) as functions of B cell lifetime. Diversity was quantified as the inverse Simpson index. Error bars indicate the standard error across 10 simulations. The rate of affinity increase was obtained from ODE simulations, whereas diversity was obtained from agent-based simulations. All simulations were performed under steady antigen supply rate (*⍺*_A_ = 125 h^−1^) and in the presence of high-sensitivity T_FH_ cells (*s* = 5).

### Influx of high-sensitivity T_FH_ cells dynamically modulates B cell selection

During the GC reaction, T_FH_ cells continuously enter existing GCs, wherein higher-sensitivity T_FH_ cells can outcompete lower-sensitivity counterparts (20, 21). To examine how such invasion events affect B cell affinity maturation, we considered a scenario in which the GC reaction was initially seeded with low-sensitivity T_FH_ cells. At day 7, under conditions of decaying antigen availability (Figure 5a), we introduced either additional low-sensitivity T_FH_ cells or high-sensitivity T_FH_ cells. High-sensitivity T_FH_ cells rapidly outcompeted resident low-sensitivity cells, ultimately dominating the T_FH_ compartment and reaching higher steady-state numbers (Figure 5b, solid curves), compared to the control case with only low-sensitivity T_FH_ cells (Figure 5b, dotted curves). Correspondingly, the entry of high-sensitivity T_FH_ cells supported a larger B cell population (Figure 5c). This takeover was accompanied by a decrease in the rate of affinity maturation (Figure 5d) and an increase in B cell diversity (Figure 5e). Together, these results suggest that the progressive influx of high-sensitivity T_FH_ cells can shift the GC reaction from an early phase characterized by rapid affinity maturation and limited diversity to a later phase with a slower affinity increase and higher diversity.

**Figure 5.**
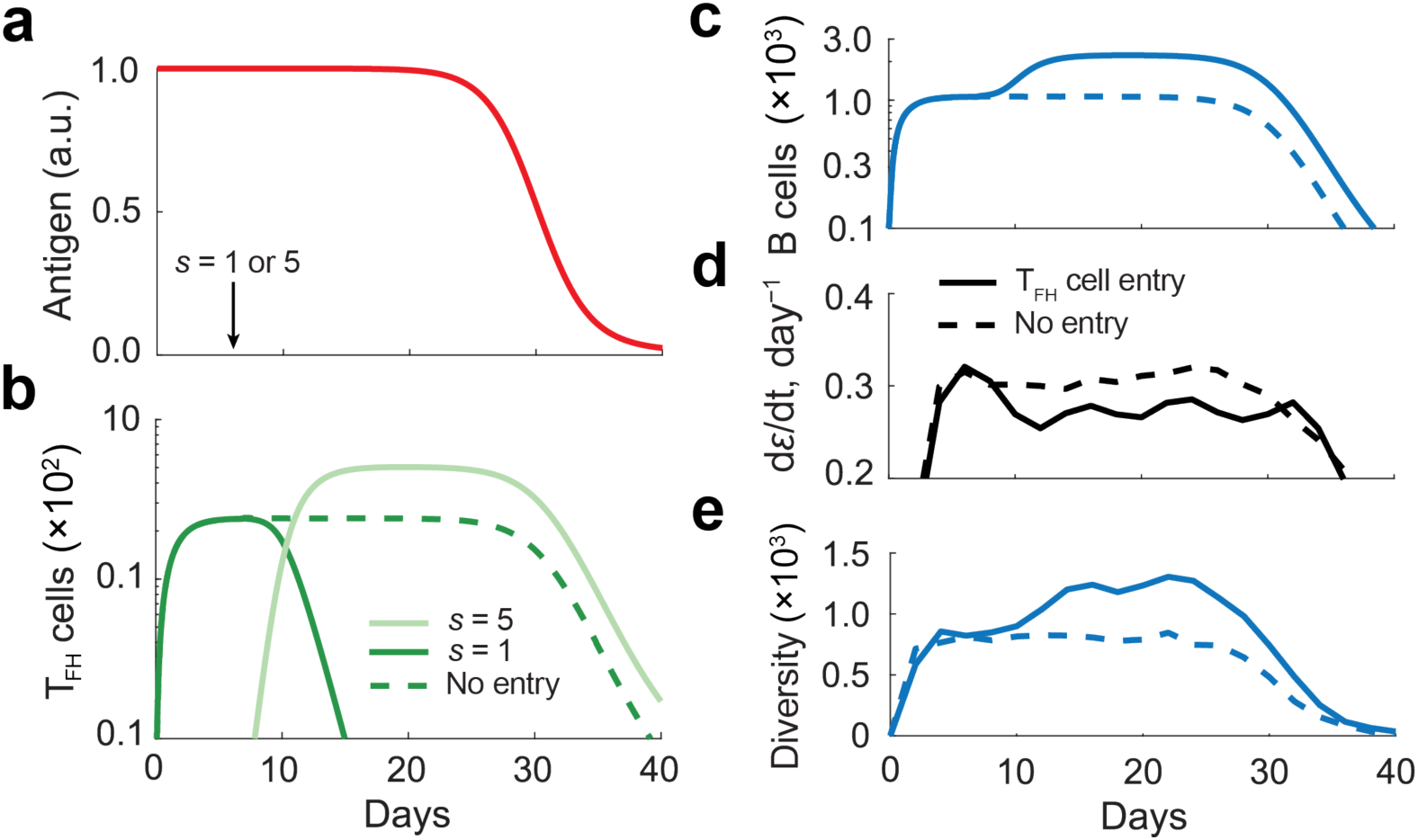
Germinal center (GC) dynamics with T_FH_ cell entry. **a)** Time-dependent antigen availability used in the simulations. Simulations were initiated with three low-sensitivity T_FH_ cells (*s* = 1); on day 7, either three additional low-sensitivity T_FH_ cells (*s* = 1) or high-sensitivity T_FH_ cells (*s* = 5) were introduced. **b, c)** Dynamics of T_FH_ cells **(b)** and B cells **(c)** when high-sensitivity T_FH_ cells were absent (dashed curves) or introduced (solid curves). **d,e)** Rate of affinity change **(d)** and B cell diversity **(e)** under the same conditions as in panels b and c. Reported values represent averages over five independent simulations, with standard errors below 10%. Diversity was quantified as the inverse Simpson index. ODE simulations were used for panels b-d whereas agent-based simulations were used to determine diversity in panel e.

## Discussion

With increasing recognition of the central role of T_FH_ cells in shaping GC dynamics, it has become clear that the collaborative interactions between T_FH_ and B cells are critical determinants of antibody quality (11, 38). Our results suggest that the balance between GC lymphocyte populations—the relative abundance and properties of B and T_FH_ cells—plays a pivotal role governing the outcome of affinity maturation. To understand how the quality and quantity of T_FH_ cells influence B cell affinity maturation, we developed a mechanistically grounded yet tractable population-dynamics model of the GC reaction that explicitly incorporates the co-evolution of B and T_FH_ cell populations. Using this theoretical framework, we found that reciprocal stimulation between B and T_FH_ cells robustly maintains the proportion of B and T_FH_ cells within the GC. While our model omits certain biological details, such as spatial organization within the GC, it captures the essential feedback mechanisms governing GC population dynamics. More detailed models that include reciprocal B-T_FH_ stimulation would therefore yield the same key conclusions, as the core regulatory logic arises from this reciprocal coupling.

Importantly, our model revealed that T_FH_ cell sensitivity tunes the balance between the speed and diversity of B cell affinity maturation by modulating the competitive landscape for T cell help. Specifically, higher-sensitivity T_FH_ cells support a broader B cell repertoire but a slower rate of affinity increase. Recent experimental findings suggest that T_FH_ cell sensitivity progressively increases during the GC reaction (20, 21). Our results provide a possible functional explanation for this observation: an early phase dominated by low-sensitivity T_FH_ cells enables rapid affinity improvement necessary for immunogen neutralization, while a later phase enriched with high-sensitivity T_FH_ cells promotes diversification of the antibody repertoire to seed memory. This process likely works in concert with the continuous influx of B cells into ongoing GC reactions, modifying the competitive landscape and further shaping the evolving B cell repertoire (39, 40).

We also found that T_FH_ cell-mediated enhancement of B cell survival, for example through secretion of cytokines such as IL-4 and IL-21, similarly shifts GC dynamics toward greater diversity at the cost of slower maturation. This trade-off may facilitate broader exploration of antibody sequence space, which is essential for discovering rare mutational trajectories leading to broadly neutralizing antibodies (bnAbs) (41–44). Indeed, recent studies such as Havenar et al. have reported that the generation of neutralizing antibodies correlates with a larger GC B cell population and a higher B:T_FH_ ratio (13). Our results provide a mechanistic basis for this link: T_FH_ cells that enhance B cell survival through cytokine support promote greater repertoire diversity, thereby increasing the likelihood of generating rare mutations necessary for bnAb formation.

Beyond T_FH_ cells, T follicular regulatory cells have also been shown to increase in proportion during later stages of the GC reaction (21). In addition, growing evidence suggests that T_FH_ cells influence B cell fate decisions, including differentiation into plasma and memory cells (11, 38, 45), highlighting that the dynamic features of the T_FH_ cell population may play a key, yet underexplored, role in long-term humoral immunity. The theoretical framework developed here offers a foundation for future studies to explore these processes, and to investigate how coordinated regulation of T and B cell populations shapes the balance between rapid protection and durable antibody diversity.

More broadly, regulation through reciprocal interactions is not unique to the GC but reflects a more general biological principle found across the body. In the gut, dynamic feedback between the microbiome and the host’s immune system shapes immune development, maintains tissue homeostasis, and enables the immune system to tolerate commensals while remaining poised to defend against pathogens (46). In the brain, the concept of the tripartite synapse highlights how astrocytes and neurons engage in bidirectional communication—astrocytes not only respond to synaptic activity but actively regulate synaptic transmission—introducing the view that brain function emerges from coordinated interactions between neurons and glia (47). These examples underscore how reciprocal feedback allows for robust, adaptive regulation in complex systems. Yet, when viewed more broadly across biological contexts, mutualisms in many co-evolving systems are unstable (48): under environmental stress, obligate cross-feeding *E. coli* populations are driven to extinction unless rescued by mutations that restore metabolic autonomy (49), while long-standing symbioses such as those between figs and fig wasps (50) or mycorrhizal fungi and plants (51) have given rise to parasitic taxa. Understanding when and how reciprocal interactions confer stability, and under what conditions they collapse or shift evolutionary character, remains a central and open question across fields—from immunology to ecology.

From a practical perspective, the concept of homeostatic reciprocal stimulation between B cells and T_FH_ cells in the GC provides new levers for tuning immune responses through vaccination. Factors that influence T_FH_ cell dynamics, such as antigen form, dose, or adjuvants, could be strategically manipulated to reshape GC architecture and enhance antibody responses. For instance, modulating T_FH_ cell sensitivity or their capacity to support B cell survival could bias selection toward high-affinity antibodies, or promote the generation of diverse repertoires to facilitate memory formation and the evolution of rare neutralizing antibodies. Understanding how the kinetics of T cell help shape B cell evolution thus opens the door to rational vaccine design strategies that guide the immune system toward specific protective outcomes.

## Materials and Methods

### Simulation Details

Simulations of Equations 2 & 3 were performed using an Euler-Maruyama integration scheme with a timestep of Δ*t* = 10^23^ h. At each timestep, population fluxes due to birth and death events were computed for both B and T_FH_ cell populations, while mutational fluxes were calculated from the birth flux of B cells, with a mutation probability per division *p_n_* given by

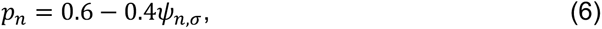

where the probability of acquiring a mutation per division depends on the magnitude of T cell help *ψ_n_*,*_ᓂ_* ∈ [0,1], resulting in mutation probabilities ranging from 0.6 for minimal help to 0.2 for maximal help (27). Unless otherwise specified, simulations were initialized with 100 B cells of identical affinity and 6 T_FH_ cells of a single sensitivity.

For the hybrid agent-based model, the same population dynamics framework was used with a timestep of Δ*t* = 3 × 10^23^ h, but each B cell’s mutation history was tracked individually. The mutation profile of each B cell was represented as a vector whose length corresponded to the total number of mutations acquired by that cell, with the *n*th entry indicating the identity of the *n*th mutations. For simplicity, all mutations were assumed to be unique.

### Mice

C57BL/6J and C57BL/6J Foxp3-GFP mice were housed at a temperature of 72 °F and humidity of 30–70% in a 12-h light/dark cycle with *ad libitum* access to food and water. Male and female mice aged 8–10 weeks at the start of the experiment were used throughout. Given the nature of the comparisons, mice were not randomized into each experimental group and investigators were not blinded to group allocation.

### Ethical Statement

All procedures in mice were performed in accordance with protocols approved by the Rockefeller University IACUC. All animal experiments were performed according to the protocols approved by the Institutional Animal Care and Use Committee of NIAID, NIH.

### Immunizations and treatments

Mice (6–12 weeks old) were immunized with 10μg or 20μg of NP17–OVA (Biosearch Technologies) precipitated in alhydrogel adjuvant (invivogen) 2% and administered into the hind footpads. Popliteal lymph nodes were then harvested at the indicated timepoints post immunization.

### Cell isolation

Single cell suspensions were prepared from lymph nodes by mechanical disruption to and were stained for surface markers in the presence of FC block. For C57BL/6J mice, intracellular Foxp3 staining was performed using the Foxp3/Transcription Factor Staining Buffer Set (eBioscience) according to the manufacturer’s instructions. For C57BL/6J Foxp3-GFP reporter mice, Foxp3 expression was assessed directly based on endogenous GFP signal following surface staining.

### Flow cytometry

Single-cell suspensions were stained with antibodies directly conjugated to surface markers. Multi-color cytometry was performed on an A3, and A5 Symphony flow cytometers (BD biosciences) and analyzed with FlowJo v10.

## Acknowledgments

This work was supported by the Natural Sciences and Engineering Research Council of Canada and a Stanford Science Fellowship (A.G.T.P.), and by Princeton University through the Center for the Physics of Biological Function. J.M. is a Branco Weiss Fellow. This project was made possible in part by grant DAF2024-342781 from the Chan Zuckerberg Initiative DAF, an advised fund of the Silicon Valley Community Foundation (N.S.W.). Additional support was provided by NIH NIGMS Grant No. R35GM146949 (B.H.G). B.H.G. is an investigator of the Chan Zuckerberg Biohub–San Francisco.

