## Supplemental information for "Modeling the Synergetic Dynamics of B cells and TFH cells in Germinal Center Reactions"

**4 Model parameters** **6**

### 1 Model

In order to investigate the influence of $T_{FH}$ cells on B cell affinity maturation, we developed a population dynamics model. The model describes B cell affinity maturation as an evolutionary hill-climbing process, wherein mutation and selection cause germinal center (GC) B cells to progressively improve their affinity. We divide the B cells into affinity classes, indexed by $n$, with $B_n$ denoting the number of B cells in class $n \in \mathbb{Z}$. Without loss of generality, we order classes so that larger $n$ corresponds to higher affinity. Within each class, a subset of B cells present antigen; these are denoted $\mathscr{B}_n$.

B cell expansion requires two steps: antigen presentation and receipt of help from T follicular helper ($T_{FH}$) cells. We denote by $T_\sigma$ the population of $T_{FH}$ cells of clone $\sigma$, and define the total $T_{FH}$ and B cell populations as

$$T = \sum_\sigma T_\sigma, \tag{S1a}$$

$$B = \sum_n B_n. \tag{S1b}$$

A B cell in affinity class $n$ that presents antigen and encounters a $T_{FH}$ cell of clone $\sigma$ proliferates at a rate proportional to the help signal strength $\psi_{n,\sigma}$. Upon proliferation, B cells may mutate: beneficial mutations increase affinity (moving up a class, $n \to n+1$), while deleterious mutations reduce affinity (moving down a class, $n \to n-1$).

Accordingly, the dynamics of B cells are given by

$$\frac{\mathrm{d}B_n}{\mathrm{d}t} = J_n - \delta_\mathrm{B} B_n + \kappa \mathscr{B}_n \sum_\sigma \psi_{n,\sigma} \frac{T_\sigma}{T+B}, \tag{S2}$$

where $J_n$ represents the net flux of B cells into class $n$ due to mutations, the second term accounts for apoptosis at rate $\delta_\mathrm{B}$, and the final term captures proliferation driven by $T_{FH}$ help, with maximum rate $\kappa$. The normalization factor $(T+B)^{-1}$ accounts for spatial competition, so that $T_\sigma/(T+B)$ represents the probability that a B cell encounters a $T_{FH}$ cell of clone $\sigma$.

$T_{FH}$ cells themselves exit the GC or undergo apoptosis at rate $\delta_\mathrm{T}$, and proliferate in proportion to productive interactions with antigen-presenting B cells. Only a fraction $\gamma$ of such interactions lead to proliferation, giving

$$\frac{\mathrm{d}T_\sigma}{\mathrm{d}t} = -\delta_\mathrm{T} T_\sigma + \gamma\kappa T_\sigma \sum_n \psi_{n,\sigma} \frac{\mathscr{B}_n}{T+B}. \tag{S3}$$

Since only antigen-presenting B cells can be selected, we also model antigen acquisition. B cells that have not yet acquired antigen do so by extracting it from follicular dendritic cells (FDCs) at rate $\beta$, weighted by the probability of successful extraction $\tilde{f}_n$. The corresponding dynamics are

$$\frac{\mathrm{d}\mathscr{B}_n}{\mathrm{d}t} = -\delta_\mathrm{B} \mathscr{B}_n + \beta \tilde{f}_n \left(B_n - \mathscr{B}_n\right) A, \tag{S4}$$

where $A$ denotes the amount of available antigen. For simplicity, our coarse-grained model neglects the loss of antigen-presenting B cells due to proliferation. This omission would effectively act as an additional death rate on the timescale of selection and division, but this contribution is subdominant to the baseline apoptosis rate, $\delta_\mathrm{B}$.

Finally, we assume that antigen is supplied to FDCs at rate $\alpha_\mathrm{A}$ and depleted only when extracted by B cells. Thus,

$$\frac{\mathrm{d}A}{\mathrm{d}t} = \alpha_\mathrm{A} - \beta A \sum_n \tilde{f}_n \left(B_n - \mathscr{B}_n\right). \tag{S5}$$

Here, the units of $A$ are defined so that one unit corresponds to the typical amount of antigen extracted by a B cell.

### 1.1 Antigen capture

In the model, B cells acquire antigen presented on FDCs in an affinity class dependent manner so that B cells expressing higher-affinity B cell receptors (BCRs) have a higher chance of acquiring antigen. For simplicity, we take this probability to be a sigmoidal function in the form

$$\tilde{f}_n = \frac{1}{1 + e^{-\varepsilon(n-\bar{n})}}, \tag{S6}$$

where $\varepsilon$ can be interpreted as the affinity difference between adjacent affinity classes, and $\bar{n}$ is the average affinity class of the B cell population.

### 1.2 $T_{FH}$ cell help signal

B cell proliferation is driven by the magnitude of help signals from $T_{FH}$ cells. In the model, the extent of help that a $T_{FH}$ cell in clone $\sigma$ provides to a B cell in affinity class $n$ is captured by $\psi_{n,\sigma}$ (see Eqs. S2 & S3).

$T_{FH}$ cells provide help in proportion to the density of antigen-derived peptide–MHC complexes (pMHCs) displayed on B cells. We assume that the amount of pMHC presented is proportional to the antigen captured by the B cell. Since $\tilde{f}_n$ (Eq. S6) denotes the probability of successful antigen capture for B cells in affinity class $n$, the amount of antigen acquired is

$$a_n = a_{\max}\tilde{f}_n, \tag{S7}$$

where $a_{\max}$ is the maximum amount of antigen that can be presented. Without loss of generality, we measure antigen concentration in units of $a_{\max}$, such that $a_{\max} = 1$.

The propensity of a $T_{FH}$ cell to provide help also depends on its intrinsic sensitivity, denoted by $s_\sigma$ for clone $\sigma$. Since no presented antigen corresponds to no help ($\psi_{n,\sigma} = 0$), and saturating antigen corresponds to maximal help ($\psi_{n,\sigma} = 1$), we assume the following functional form:

$$\psi_{n,\sigma} = \frac{1}{1 + \frac{(1/s_\sigma)}{a_n}}. \tag{S8}$$

Here, $1/s_\sigma$ can be interpreted as the amount of antigen required for a B cell to receive 50% of the maximum help signal from $T_{FH}$ cells in clone $\sigma$.

### 1.3 Flux due to mutations

In the GC reaction, B cells undergo somatic hypermutation during proliferation. Within this model, the effects of these mutations are incorporated via the mutation flux term $J_n$ in Eq. S2. Let $p_n$ reflect the probability of mutations occurring, such that the rate at which mutants are generated from class $n$ is

$$\mu_n B_n = p_n\, \kappa\, \mathscr{B}_n \sum_\sigma \frac{\psi_{n,\sigma} T_\sigma}{T + B}. \tag{S9}$$

Recent studies have shown that B cells receiving stronger T cell help undergo fewer mutations upon division [1, 2]. To capture this dependence, we model the mutation probability per division, $p_n$, as

$$p_n = p_{\max} - (p_{\max} - p_{\min})\, \bar{\psi}_n, \tag{S10}$$

where $\bar{\psi}_n = (1/T)\sum_\sigma T_\sigma \psi_{n,\sigma}$ represents the average T cell help received by B cells of class $n$. Here, $p_{\min}$ and $p_{\max}$ denote the minimum and maximum mutation probabilities, corresponding respectively to the cases of maximal help ($\bar{\psi}_n = 1$) and minimal help ($\bar{\psi}_n = 0$).

Mutations can be beneficial, deleterious, silent, or lethal. For simplicity, we only account for silent, beneficial and deleterious mutations, which occur with probabilities $p_0$, $p_+$ and $p_-$, respectively, conditional on a mutation event. Accordingly, we divide the mutant flux (Eq. S9) into contributions from beneficial and deleterious mutants as

$$J_{\chi,n} = p_\chi \mu_n B_n, \tag{S11}$$

where $\chi \in \{0, +, -\}$, and $J_{0,n}$, $J_{+,n}$ and $J_{-,n}$ denote, respectively, the flux of silent, beneficial and deleterious mutants generated from B cells in affinity class $n$. The net flux of B cells into affinity class $n$ due to mutations is then

$$J_n = -(J_{+,n} + J_{-,n}) + J_{+,n-1} + J_{-,n+1}, \tag{S12}$$

so that, by construction, $\sum_n J_n = 0$.

### 2 Fixed-point analysis

To build intuition, we first study a simplified version of the model. We neglect B cell affinity classes (so no mutation flux) and assume the $T_{FH}$ population is monoclonal. In this limit, the dynamics reduce to

$$\frac{dB}{dt} = -\delta_B B + \kappa\psi \frac{T\mathscr{B}}{T+B}, \tag{S13a}$$

$$\frac{dT}{dt} = -\delta_T T + \gamma\kappa\psi \frac{T\mathscr{B}}{T+B}, \tag{S13b}$$

$$\frac{d\mathscr{B}}{dt} = -\delta_B \mathscr{B} + \tfrac{1}{2}\beta\big(B-\mathscr{B}\big)A, \tag{S13c}$$

$$\frac{dA}{dt} = \alpha_A - \tfrac{1}{2}\beta\big(B-\mathscr{B}\big)A. \tag{S13d}$$

Here, with no affinity classes, $a_n, \tilde{f}_n \to 1/2$, and the $T_{FH}$ help signal becomes constant ($\psi = s/(2+s)$), where $s$ denotes the sensitivity of the $T_{FH}$ cell clone.

#### 2.1 Quasi–steady-state reduction

To simplify the analysis, we assume antigen dynamics are fast compared to lymphocyte dynamics. Thus, antigen capture and presentation rapidly equilibrate. In this quasi–steady-state limit,

$$A = \frac{2\alpha_A}{\beta(B-\mathscr{B})}, \tag{S14}$$

which implies

$$\mathscr{B} = \frac{\alpha_A}{\delta_B}. \tag{S15}$$

Substituting this back into the dynamics (Eqs. S13), the lymphocyte equations reduce to

$$\frac{dB}{d\tilde{t}} = -\tilde{\delta}_B B + \psi \frac{T}{T+B}, \tag{S16}$$

$$\frac{dT}{d\tilde{t}} = -\tilde{\delta}_T T + \gamma\psi \frac{T}{T+B}, \tag{S17}$$

where tildes represent rescaled variables with time in units of $\delta_B/(\kappa\alpha_A)$.

It follows from Eqs. S16 & S17 that this system admits a non-trivial fixed point given by

$$B^* = \frac{\gamma\psi}{\gamma\tilde{\delta}_B + \tilde{\delta}_T}, \qquad T^* = \frac{\gamma^2\psi\,\tilde{\delta}_B}{\tilde{\delta}_T(\gamma\tilde{\delta}_B + \tilde{\delta}_T)}. \tag{S18}$$

In particular, the total abundance of lymphocytes is given by a simple expression,

$$T^* + B^* = \frac{\gamma\psi}{\tilde{\delta}_T} = \frac{\gamma\psi}{\delta_T}\left(\frac{\kappa\alpha_A}{\delta_B}\right), \tag{S19}$$

which is proportional to the antigen supply rate $\alpha_A$, so that higher antigen availability leads to a larger lymphocyte population.

#### 2.2 Stability of lymphocyte population

The Jacobian of the lymphocyte dynamics (Eqs. S16 & S17) is

$$\mathcal{J}(B,T) = \begin{bmatrix} -\tilde{\delta}_{\mathrm{B}} - \psi\frac{T}{(T+B)^2} & \psi\frac{B}{(T+B)^2} \\ -\gamma\psi\frac{T}{(T+B)^2} & -\tilde{\delta}_{\mathrm{T}} + \gamma\psi\frac{B}{(T+B)^2} \end{bmatrix}. \tag{S20}$$

At the fixed point, the trace and determinant of the Jacobian are

$$\operatorname{tr}\mathcal{J}(B^*,T^*) = -\frac{\tilde{\delta}_{\mathrm{B}}}{\gamma\tilde{\delta}_{\mathrm{B}} + \tilde{\delta}_{\mathrm{T}}}\left[\gamma\tilde{\delta}_{\mathrm{B}} + (2+\gamma)\tilde{\delta}_{\mathrm{T}}\right] < 0, \tag{S21}$$

$$\det\mathcal{J}(B^*,T^*) = \tilde{\delta}_{\mathrm{B}}\,\tilde{\delta}_{\mathrm{T}} > 0. \tag{S22}$$

Since the death rates $\tilde{\delta}_{\mathrm{B/T}}$ and $\gamma$ are both positive, the trace is negative and the determinant is positive. Therefore, the fixed point $(B^*, T^*)$ (Eq. S18) is linearly stable.

#### 2.3 B:T$_{\mathrm{FH}}$ ratio

From Eq. S18, it follows that the steady-state B:T$_{\mathrm{FH}}$ ratio $r^*$ reduces to a simple expression

$$r^* \equiv \frac{B^*}{T^*} = \frac{\delta_{\mathrm{T}}}{\gamma\delta_{\mathrm{B}}}. \tag{S23}$$

Note that the B:T$_{\mathrm{FH}}$ ratio only depends on the death rates and $\gamma$, and is independent of T$_{\mathrm{FH}}$ cell sensitivity or antigen availability.

### 3 Reduced model

For tractability, we focus on the case where antigen capture dynamics are much faster than the population-level dynamics of lymphocytes. This separation of timescales allows us to treat Eqs. S4 & S5 as being in quasi-steady state. In this limit, the coupled dynamics of B cells and antigen are described by

$$\mathscr{B}_n^* = \frac{\beta\tilde{f}_n A^*}{\delta_{\mathrm{B}} + \beta\tilde{f}_n A^*} B_n, \tag{S24}$$

$$A^* = \frac{\alpha_{\mathrm{A}}}{\beta\sum_n \tilde{f}_n(B_n - \mathscr{B}_n^*)}. \tag{S25}$$

Combining these relations yields a self-consistency equation for the number of antigen-presenting B cells given by

$$\mathscr{B}_n^* = \frac{\alpha_{\mathrm{A}}\tilde{f}_n}{\delta_{\mathrm{B}}\sum_m f_m(B_m - \mathscr{B}_m^*) + \alpha_{\mathrm{A}}\tilde{f}_n} B_n. \tag{S26}$$

This nonlinear relation admits analytic solutions only when the number of affinity classes is three or fewer (which typically, won't be the case). For larger numbers of affinity classes, the equation reduces to a polynomial of degree five or higher, for which, by the Abel–Ruffini theorem, no general solution in closed form exists.

#### 3.1 Reduced lymphocyte dynamics in the high-competition limit

We assume strong competition for antigen, such that $\mathscr{B}_n^*/B_n \ll 1$ due to antigen limitation. Under this assumption, the steady-state fraction of antigen-presenting B cells simplifies to

$$\mathscr{B}_n^* \approx \frac{\alpha_{\mathrm{A}}\tilde{f}_n B_n}{\delta_{\mathrm{B}}\sum_m f_m B_m}. \tag{S27}$$

Using the quasi-static approximation for antigen, the B cell and $T_{FH}$ cell dynamics are

$$\frac{\mathrm{d}B_n}{\mathrm{d}\tilde{t}} = \tilde{J}_n - \tilde{\delta}_B B_n + \sum_{\sigma} B_n \Psi_{n,\sigma}(\vec{B}, \vec{T}) T_\sigma, \tag{S28}$$

$$\frac{\mathrm{d}T_\sigma}{\mathrm{d}\tilde{t}} = -\tilde{\delta}_T T_\sigma + \gamma \sum_{n} B_n \Psi_{n,\sigma}(\vec{B}, \vec{T}) T_\sigma, \tag{S29}$$

where the rescaled time $\tilde{t}$ is measured in units of $1/\kappa$, and the effective interaction term is

$$\Psi_{n,\sigma}(\vec{B}, \vec{T}) T_\sigma = b(\alpha_A) f_n \frac{T_\sigma}{T + B} \psi_{n,\sigma}, \tag{S30}$$

where $b(\alpha_A) = (\alpha_A/\delta_B)/B$ is the fraction of B cells presenting antigen, and $f_n = \tilde{f}_n/\langle \tilde{f}_n \rangle$ with $\langle \tilde{f}_n \rangle = \sum_n B_n \tilde{f}_n / B$ quantifies the ability of different B cell classes at capturing antigen. This effective interaction $\Psi_{n,\sigma}$ can be interpreted as the product of two contributions: (i) a modulating factor $b(\alpha_A) f_n$, which represents the fraction of B cells in affinity class $n$ that have captured antigen, and (ii) an intrinsic interaction $T_\sigma \psi_{n,\sigma}/(T + B)$, which accounts for sensitivity and availability (e.g. due to steric interference) depending on the lymphocyte populations and composition of $T_{FH}$ cells.

#### 3.2 Steady-state population and B:$T_{FH}$ ratio

Summing over all B cell affinity classes in Eq. S28 and all T cell clones in Eq. S29 gives the population-level dynamics

$$\frac{\mathrm{d}B}{\mathrm{d}\tilde{t}} = \sum_n \tilde{J}_n - \tilde{\delta}_B B + \sum_{n,\sigma} B_n \Psi_{n,\sigma}(\vec{B}, \vec{T}) T_\sigma, \tag{S31}$$

$$\frac{\mathrm{d}T}{\mathrm{d}\tilde{t}} = -\tilde{\delta}_T T + \gamma \sum_{n,\sigma} B_n \Psi_{n,\sigma}(\vec{B}, \vec{T}) T_\sigma. \tag{S32}$$

At steady state, the B:$T_{FH}$ ratio is

$$r^* = \frac{\delta_T}{\gamma \delta_B}, \tag{S33}$$

which coincides with the case without explicit B cell affinity classes (Eq. S23).

### 4 Model parameters

Most parameters in the model were determined directly from experimental measurements, while a few were inferred from related quantities. In particular, we selected the maximum B-cell proliferation rate, $\kappa$, such that the GC reaches a steady state within a few days. The relative proliferation factor, $\gamma$, was selected to produce a GC containing a biologically reasonable number of B cells.

| Parameter | Description | Value | Reference |
|---|---|---|---|
| $1/\delta_B$ | B cell lifetime | 8 h, variable | [3] |
| $1/\delta_T$ | $T_{FH}$ cell lifetime | 18 h | [4] |
| $\varepsilon$ | Binding energy difference between affinity classes | 0.5 $k_B T$ | [6] |
| $1/\kappa$ | Effective B cell proliferation timescale | 2 h | N/A |
| $\gamma$ | Relative proliferation factor | 0.1 | N/A |

Table 1: Model parameters used in simulations.

Mutation parameters were set to $p_{\min} = 0.2$ and $p_{\max} = 0.6$ [1]. Upon mutation, the probabilities of affinity-affecting mutations were $p_+ = 0.19$ and $p_- = 0.01$ [6].
